# Extensive transcriptional changes in the aphid species *Myzus cerasi* under different host environments associated with detoxification genes

**DOI:** 10.1101/366450

**Authors:** Peter Thorpe, Carmen M. Escudero-Martinez, Sebastian Eves-van den Akker, Jorunn I.B. Bos

## Abstract

Aphids are phloem-feeding insects that cause yield losses to crops globally. These insects feature complex life cycles, which in the case of many agriculturally important species involves the use of primary and secondary host plant species. Whilst host alternation between primary and secondary host can occur in the field depending on host availability and the environment, aphid populations maintained as laboratory stocks generally are kept under conditions that allow asexual reproduction by parthenogenesis on secondary hosts. Here, we used *Myzus cerasi* (black cherry aphid) to assess aphid transcriptional differences between populations collected from primary hosts in the field and those adapted to secondary hosts under controlled environment conditions. Adaptation experiments of *M. cerasi* collected from local cherry tress to reported secondary host species resulted in low survival rates. Moreover, aphids were unable to survive on secondary host Land cress, unless first adapted to another secondary host, cleavers. Transcriptome analyses of populations collected from primary host cherry in the field and the two secondary host plant species in a controlled environment showed extensive transcriptional plasticity to a change in host environment, with predominantly genes involved in redox reactions differentially regulated. Most of the differentially expressed genes across the *M. cerasi* populations from the different host environments were duplicated and we found evidence for differential exon usage. In contrast, we observed only limited transcriptional to a change in secondary host plant species.

## Introduction

Aphids are phloem-feeding insects that belong to the order Hemiptera. Insects within this order feature distinctive mouthparts, or stylets, that in the case of phytophagous insects, are used to pierce plant tissues and obtain nutrients from the plant phloem. One striking feature of the complex life cycle of about 10 % of aphid species is the seasonal host switching between unrelated primary (winter) and secondary (summer) host plants, also called host alternation or heteroecy (Mordvilko 1928); (Williams 2007). Host alternating aphids predominantly use woody plants as their primary hosts, on which (overwintering) eggs are laid, from which the first generation of aphids, or fundatrices, emerge in spring. The fundatrices, and their offspring, reproduce by parthenogenesis (asexual reproduction), giving birth to live nymphs. Winged forms (alate) will migrate to secondary host plants over the summer months where the aphid populations will go through multiple parthenogenic generations. In autumn, sexual female and male aphids will reproduce sexually and overwintering eggs are laid on the primary host. Exceptions to this general life cycle exist, with some aphids for example having multi-year cycles (Kennedy 1959).

Heteroecy in aphids has independently arisen in different aphid lineages throughout evolutionary history (Moran 1988) with monoecy (with the entire life cycle taking place on one plant species) on trees thought to be the ancestral state. Many different hypotheses explain the maintenance of heteroecy and driving factors described include nutritional optimization, oviposition sites, natural enemies, temperature tolerance, and fundatrix specialization (Moran 1988). It is likely that switching between host plant species requires aphids to adapt to differences in host nutritional status as well as potential differences in plant defense mechanisms against insects. Host plant specialization in the pea aphid species complex is associated with differences in genomic regions encompassing predicted salivary genes as well as olfactory receptors (Jaquiéry et al. 2012). Moreover, adaptation of *M. persicae* to different secondary host plant species involves gene expression changes, including of genes predicted to encode for cuticular proteins, cathepsin B protease, UDP-glycosyltransferases and P450 monooxygenases (Mathers et al. 2017). Aphid secondary hosts include many important agricultural crops and are generally more suitable for maintaining clonal (asexual) aphid laboratory stocks used for research experiments. To what extend aphid gene expression is affected upon collecting aphids from the field and adapting them to select secondary host plants in a laboratory environment remains unclear.

*Myzus cerasi*, or black cherry aphid, uses mainly *Prunus cerasus* (Morello cherry) and *Prunus avium (*sweet cherry), but also other *Prunus* species as primary hosts and several herbaceous plants (*Gallium* spp., *Veronica* spp., and cruciferous species) as secondary hosts (Barbagallo et al. 2017) (Blackman and Eastop 2000). Infestation can cause significant damage on cherry trees, due to leaf curling, and shoot deformation, pseudogall formation, as well as fruit damage. Recently, we generated a draft genome for *M. cerasi*, providing novel insights into potential parasitism genes as well as genome evolution (Thorpe et al. 2018). The increasing availability of genomics resources for aphids, including *M. cerasi*, facilitates further understanding of aphid biology, including the processes involved adapting to a change in host environment. In this study, we collected *M. cerasi* from primary hosts (cherry trees) in the field with the aim to adapt these to different reported secondary hosts, *Galium aparine* (cleavers) and *Barbarea verna* (Land cress) under laboratory conditions for comparative transcriptome analyses. We found that aphids collected from their primary host in the field differed in their ability to adapt to the secondary host plant species in a controlled environment, with no aphids surviving transfer to *Barbarea verna* (Land cress). We compared the transcriptomes of *M. cerasi* aphids adapted under laboratory conditions to secondary hosts *Galium aparine* (cleavers) and *Barbarea verna* (Land cress) and only observed limited transcriptional changes. However, when comparing the transcriptomes of these adapted aphids to field collected aphids from primary hosts, we noted extensive transcriptional changes, especially with regards to predicted detoxification genes. The majority of differentially expressed genes were duplicated, implicating multigene families in aphid adaptation to host environments.

## Results and discussion

### *Myzus cerasi* host adaptation under controlled laboratory conditions is associated with low survival rates

When attempting to establish a colony of *M. cerasi* from populations occurring on local cherry trees, we observed differences in survival rates upon transfer to reported secondary host plant species under controlled plant growth conditions. While aphids were unable to survive transfer from primary host cherry to Land cress (*Barbarea verna*), we observed a 10%-20% survival rate upon transfer to cleavers (*Galium aparine*) (Fig. 1). However, once aphid populations were established on cleavers, individuals from this population were successfully transferred to cress plants. We performed similar field to lab host transfer experiments with aphids collected from cherry trees at two different locations in three independent replicates with similar results (Fig. 1). Our observation that *M. cerasi* collected from primary hosts in the field shows a difference in ability to infest two reported secondary hosts likely reflects these aphids face a major hurdle represented by a change in host environment. In addition, our finding raises the question whether aphids may expand their secondary host range once adapted to a preferred secondary host species.

**Fig. 1.**
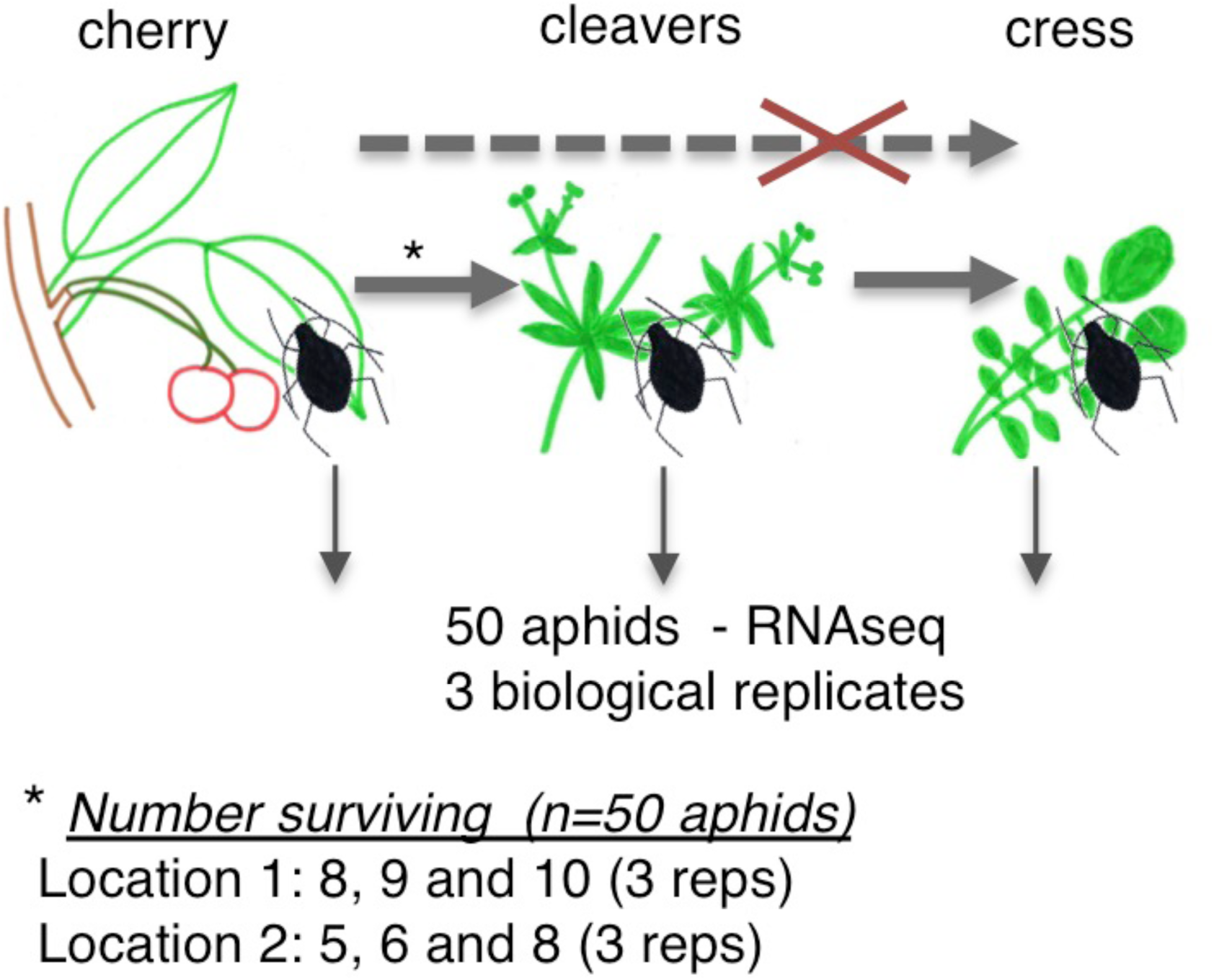
Schematic overview of host environment adaptation experiments and aphid survival rates. *Myzus cerasi* aphids were collected from cherry trees at two separate field locations. None of the aphids collected from cherry in the field were able to survive directly on *Barbarae verna* (Land cress) plants. However, a 10-20% survival rate was recorded when aphids were moved onto *Galium aparine* (cleavers) in a controlled environment. The host adaptation experiments were performed in 3 biological replicates.

### *Myzus cerasi* shows extensive transcriptional plasticity to a change in host environment

We assessed the changes that take place at the transcriptional level in *M. cerasi* when adapting the field-collected aphids from cherry to secondary hosts in a controlled plant growth environment. Specifically, we sequenced the transcriptomes of *M. cerasi* populations collected from cherry (field conditions), and of aphids established over a 3-week period on cleavers or cress (controlled environment) using RNAseq.

We performed differential gene expression analysis (LOG fold change >2, False Discovery Rate (FDR) p<0.001) between the different aphid populations to identify gene sets associated with the different host plant environments. Cluster analyses of the aphid transcriptional responses from this and previous work reporting on differential aphid gene expression in head versus body tissues (Thorpe et al. 2016) revealed that the overall expression profiles could be distinguished based on the aphid tissue used for sample preparation as well as the host environment (Fig. S1A). Indeed, principal component analyses showed a clear separation between aphid transcriptomes associated with primary host (field conditions) and the different secondary hosts (controlled conditions) (Fig. S1B). Overall, we identified 934 differentially expressed genes by comparing the different datasets for each of the aphid populations (Fig. 2A, Table S1). A heat map of these 934 genes shows that gene expression profiles from aphids adapted to secondary hosts (cleavers and cress) and maintained in a controlled environment are more similar to each other than to the gene expression profiles of aphids collected from primary hosts in the field (Fig. 2A). Co-expression analyses reveals six main clusters of differentially expressed genes, two of which (A and E) contain the majority of genes (Fig. 2A and 2B). Cluster A contains 493 genes, which show higher expression in aphids maintained on secondary hosts (controlled environment) versus those collected from primary hosts (field), and cluster B contained 342 genes showing an opposite profile. GO annotation revealed over-representation of terms associated with oxido-reductase activity in both clusters, as well as several terms associated with carotenoid/tetrapenoid biosynthesis in case of cluster B (Table S2).

**Fig. 2.**
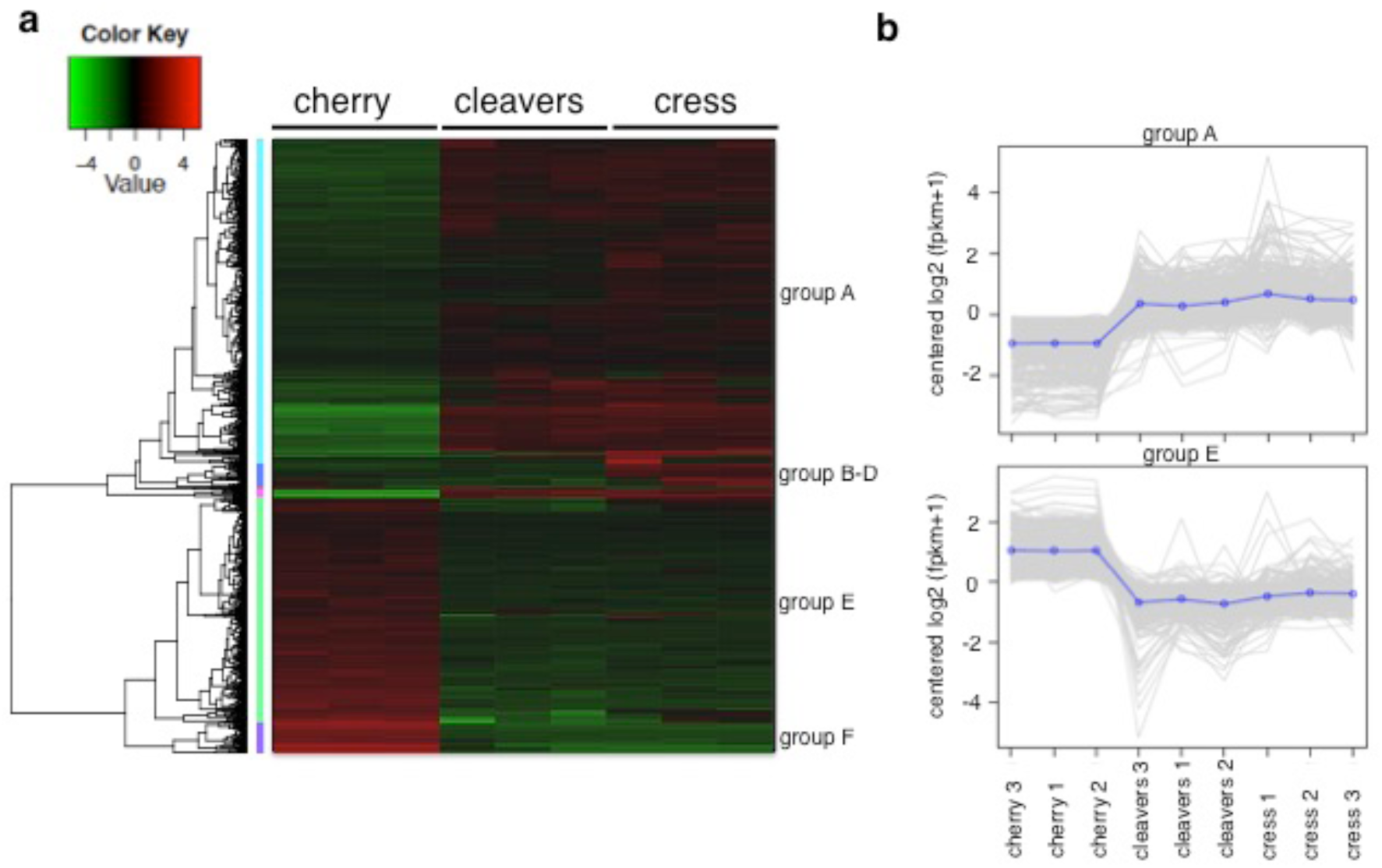
Clustering of differentially expressed genes across *Myzus cerasi* populations from primary (field) and secondary (controlled environment) hosts. (A) Cluster analyses of the 934 genes differentially expressed in *M. cerasi* populations from different host environments. (B) Expression profiles of the 493 co-regulated genes in cluster A and of the 342 coregulated genes in cluster E.

To assess differential expression of *M. cerasi* genes across the host environment-aphid interactions we also analyzed pairwise comparisons for differentially expressed gene sets. The largest set of differentially expressed genes (736) was found in comparisons between aphids from cherry (primary host, field) and cress (secondary host, controlled environment), with 443 genes more highly expressed in aphids from cress, and 293 more highly expressed in aphids from cherry (Fig. 3A). A total of 733 differentially expressed genes were found in comparisons of aphids from cherry (primary host, field) versus cleavers (secondary host, controlled environment, with 367 genes more highly expressed in aphids collected from cherry and 366 genes more highly expressed in aphids collected from cleavers (Fig. 3A). The higher number of genes up-regulated in the cress-cherry comparison may reflect the difficulties in adapting to this secondary host species, with *M. cerasi* unable to infest cress when collected from cherry (field). Our differential gene expression data not only is impacted by a change in host plant species, but also a change in plant growth environment to which the aphids were exposed. To what extend both these factors contribute the observed gene expression changes remains an open question. Nevertheless, aphids cannot make the transition to cress in the controlled environment unless they first parasitise cleavers in the controlled environment. This suggests that the transition to cleavers enables subsequent transitions through a yet unidentified mechanism.

**Fig. 3.**
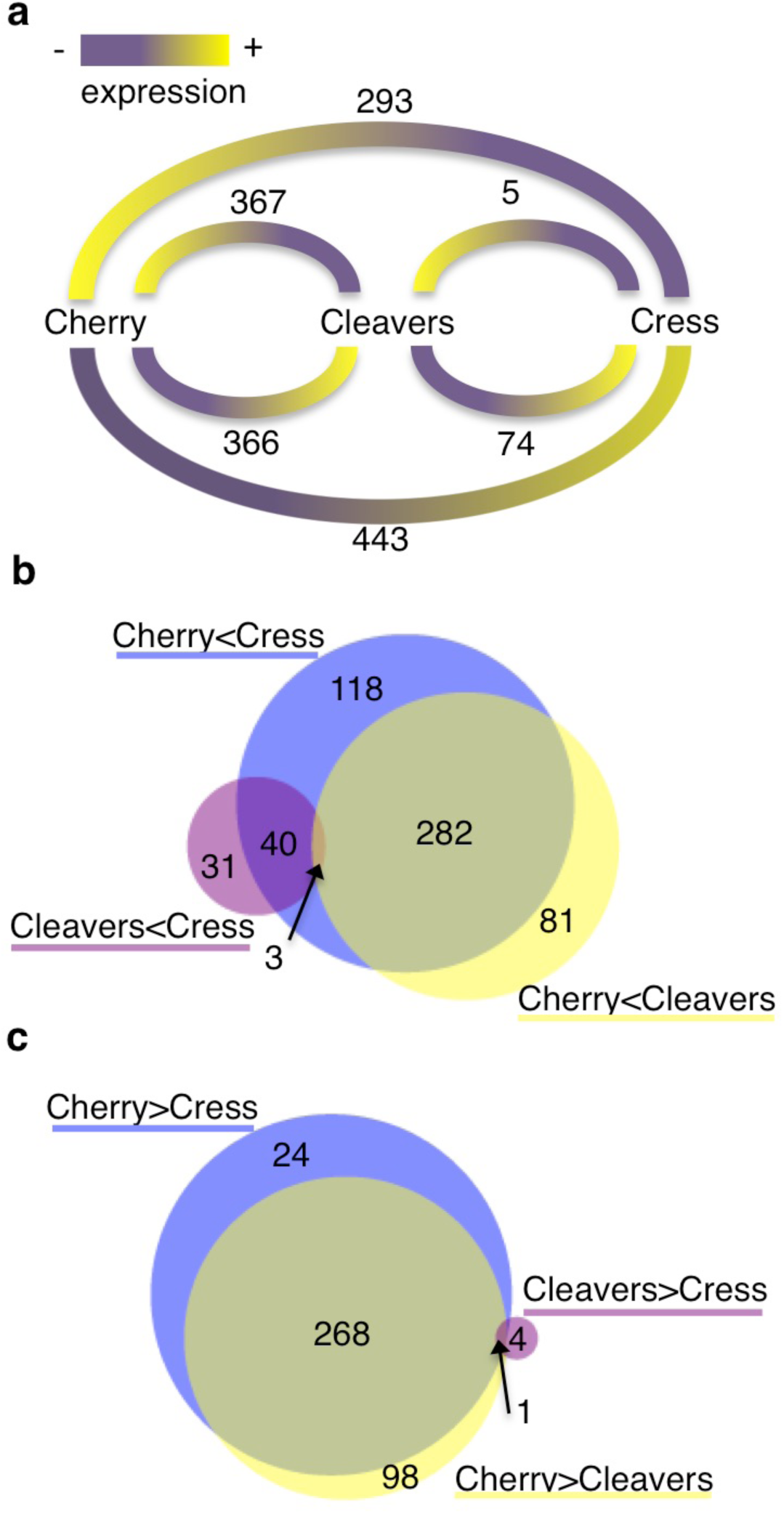
Differentially expressed genes in pairwise comparisons between the different *Myzus cerasi* populations. (A) Numbers of genes for each pairwise comparison between aphids collected form the different host species, cherry (field), cleavers (controlled environment) and cress (controlled environment). Yellow color indicates high level of expression, whereas purple color indicates low expression in the different pairwise comparisons. (B) Venn diagram showing the overlap in differentially expressed genes sets that are lower expressed in the aphids from primary host cherry (field) compared to those collected from secondary hosts cleavers and cress (both in a controlled environment), and also lower expressed in aphids from cleavers than those from cress. (C) Venn diagram showing the overlap in differentially expressed genes sets that are higher expressed in the aphids from primary host cherry (field) compared to those collected from secondary hosts cleavers and cress (both in a controlled environment), and also higher expressed in aphids from cleavers than those from cress.

A relatively small number of genes were differentially expressed between aphids collected from the two secondary hosts cleavers and cress (both grown under controlled conditions), with only 5 genes more highly expressed in aphids from cleavers, and 74 genes more highly expressed in aphids form cress (Fig. 3A). This suggests that *M. cerasi* shows limited transcriptional plasticity to a switch in secondary host environment, once adapted. This is in line with our previous observation that only a relatively small set of genes is differentially expressed in *M. persicae* and *R. padi* when exposed to different host or non-/poor-host plants (Thorpe et al. 2018) as well as the relatively small number of transcriptional changes when *M. persicae* is adapted to different secondary hosts (Mathers et al. 2017).

GO enrichment analyses of the 443 genes more highly expressed in aphids collected from cress (controlled environment) compared to those collected from cherry (field) shows overrepresentation of genes predicted to be involved various processes, including in heme binding (GO:0020037), tetrapyrrole binding (GO:0046906), monooxygenase activity (GO:0004497), oxidoreductase activity (GO:0016705), iron ion binding (GO:0005506), and hydrolase activity (GO:0016787) (Table S3). This set of 443 genes contains 282 of the 366 genes that are also more highly expressed in aphids from the other secondary host plant species, cleavers, with similar GO annotations (Fig. 3B; Table S4). The 293 genes more highly expressed in aphids collected from cherry (field) than those from cress (controlled environment) shows over-representation of genes predicted in oxidoreductase activity (GO:0016620, GO:0016903, GO:0055114, GO:001649) as well as other processes such as fatty-acyl-CoA reductase (alcohol-forming) activity (GO:0080019), interspecies interaction between organisms (GO:0044419), and symbiosis (GO:0044403) (Table S3). For the gene sets differentially expressed between aphids collected from cherry (field) and cleavers (controlled environment), GO enrichment analyses reveal that in reciprocal comparisons genes predicted to function in redox reactions are also over-represented (Table S3). The over-representation of differentially expressed genes involved in redox across the different *M. cerasi* populations likely reflects different requirements for aphids under different host environment conditions.

Interestingly, among the 367 genes more highly expressed in aphids collected from cherry (field) compared to those collected from cleavers (controlled environment), we found that the majority of GO terms identified through enrichment analyses correspond to metabolic processes (Table S3). Of these 367 transcripts 268 show similar expression differences in aphids collected from cherry (field) versus those collected from cress (controlled environment), whereas 98 are specific to the comparison of aphids collected from cherry (field) versus cleavers (controlled environment) (Fig. 3C). Whilst GO enrichment analyses showed over-representation of genes involved in redox reactions in the set of 268 overlapping transcripts, the 98 transcripts specifically up-regulated in aphids collected from cherry (field) versus cleavers (controlled environment) show over-representation in metabolic processes, and especially those associated with terpenoid/carotenoid biosynthesis which are involved in aphid pigmentation (Table S5) (Moran and Jarvik 2010). Possibly this observation reflects that *M. cerasi* requires specific gene sets for pigmentation and feeding under specific host environmental conditions. Notably, we did not observe any noticeable change in aphid color upon adapting aphids from primary hosts in the field to secondary hosts in the lab. Aphids featured a dark brown to black color on all plant species tested (not shown), suggesting the differential regulation of carotenoid genes is not associated with aphid color in this case but with other unknown physiological functions.

To independently test whether select *M. cerasi* genes were differentially expressed in aphids collected from primary (field) and secondary (controlled environment) host plants, we repeated the collection of aphids from local cherry trees (separate site, location 2) and performed adaptation experiments to cleavers and cress. We selected 10 genes for independent validation of expression profiles by qRT-PCR. Five of these 10 genes were selected based on enhanced expressed in aphids from cherry (field) compared to aphids from secondary (controlled environment) host plants, and another 5 genes for being more highly expressed in aphids from secondary (controlled environment) host plants compared to aphids from cherry (field). The genes selected based on higher expression in aphids from cherry (field) showed similarity to genes predicted to encode a peroxidase, RNA-binding protein 14-like, hybrid sensor histidine kinase response regulator, maltase isoform a, and a lactase-phlorizin hydrolase. The genes selected based on higher expression in aphids from secondary (controlled environment) hosts showed similarity to genes predicted to encode an unknown protein, a venom-like protease, a thaumatin-like protein, protein kintoun, and a cytochrome P450. Except for the gene with similarity to a venom-like protease, all genes showed a similar gene expression profile in both samples used for the RNAseq experiments and in the independently collected and adapted aphids from a different site, indicating that this gene set is consistently differentially expressed when *M. cerasi* was adapted from primary hosts in the field to secondary hosts in a controlled environment (Fig. S2). Most of these genes have predicted functions in detoxification, in line with our hypothesis that aphids require different sets of genes to deal with potential defensive plant compounds associated with different host environments. To what extend our observations are associated with primary versus secondary host factors or field versus controlled environment factors is not clear. Notably, in *H. persikonus* collected from primary and secondary host plant species in the field, a similar observation was made in that an extensive gene set associated with detoxification was differentially regulated (Cui et al. 2017).

### Single Nucleotide Polymorphism (SNP) analyses suggest that an aphid sub-population is able to adapt from primary hosts in the field to secondary hosts in a controlled environment

We used the transcriptome dataset we generated here to compare the level of sequence polymorphisms between the aphid populations from the different primary (field) and secondary (controlled environment) host plants species. Variants/SNPs were predicted by mapping the RNAseq dataset for each aphid population (cherry, cleavers and cress) to the *M. cerasi* reference genome for each condition, with only unique mapping being allowed. The number of SNPs within each 10Kb window was calculated. The *M. cerasi* population from cherry has significantly more SNPs per 10Kb than the populations from both cleavers and cress when mapping reads back to the reference genome (p<0.001, Kruskal-Wallis with Bonferroni post hoc correction). In contrast, the aphid populations from cleavers and cress showed no significant difference in the number of SNPs per 10Kb (p=0.29, Kruskal-Wallis with Bonferroni post hoc correction). These results are consistent with the observation that the population of *M. cerasi* went through a bottle neck during the transfer from cherry to cleavers, but not cleavers to cress. Based on these findings we propose that only a subpopulation of the primary host population may switch to secondary host plant species. It should be noted that these data are based on RNAseq, and do not rule out the possibility of allele-specific expression across the different host interactions. Hence, further characterization of the *M. cerasi* (sub)populations using DNAseq will be required to gain further insight into adaptation of this aphid species to its hosts.

### Limited differential expression of predicted *M. cerasi* effectors across field and adapted populations

We assessed whether predicted *M. cerasi* effectors are differentially expressed in aphid population collected from a primary host (field) or secondary hosts plants (controlled environment). The 224 predicted *M. cerasi* effectors we previously identified (Thorpe et al. 2018) show a wide range of expression levels across different interactions, with most expression variation in aphids collected from cherry under field conditions (Fig. 4A; Table S6). However, when assessing expression of a random non-effector set of similar size, this expression variation in aphids was less pronounced (Fig. 4A). Despite the observed variation in expression patterns, we only found a small number of differentially expressed candidate effectors, mainly when comparing aphids collected from the primary (field) versus secondary hosts (controlled environment). Specifically, 13 candidate effectors are more highly expressed in aphids from both secondary host species (controlled environment) compared to aphids from the primary host cherry (field), with one additional candidate effector more highly expressed in the case of aphids from cleavers compared to cherry only (Mca17157|adenylate kinase 9-like) (Table S6). Although these candidate effectors were mainly of unknown function, several show similarity to thaumatin-like proteins and a venom protease. Interestingly, the candidate effector with similarity to the venom protease, Mca05785 (upregulated in secondary hosts (controlled environment)), is member of a venom protease gene family cluster that consists of four members (3 are tandem duplications, 1 is a proximal duplication). Three of these are predicted to encode secreted proteins, and all members show higher expression levels in aphids from secondary hosts under controlled conditions compared to aphids from primary host in the field, but this variation was below the LOG2 fold change cut-off (Table S6). In addition, 1 candidate effector (similar to RNA-binding protein 14) was differentially expressed when comparing aphids from the two secondary host plants (controlled environment), and 5 candidate effectors (Mca07285, Mca07514, Mca16980, Mca07516, Mca09259) were more highly expressed in aphids collected from cherry (field) compared to aphids from cleavers and/or cress (controlled environment) (Table S6).

**Fig. 4.**
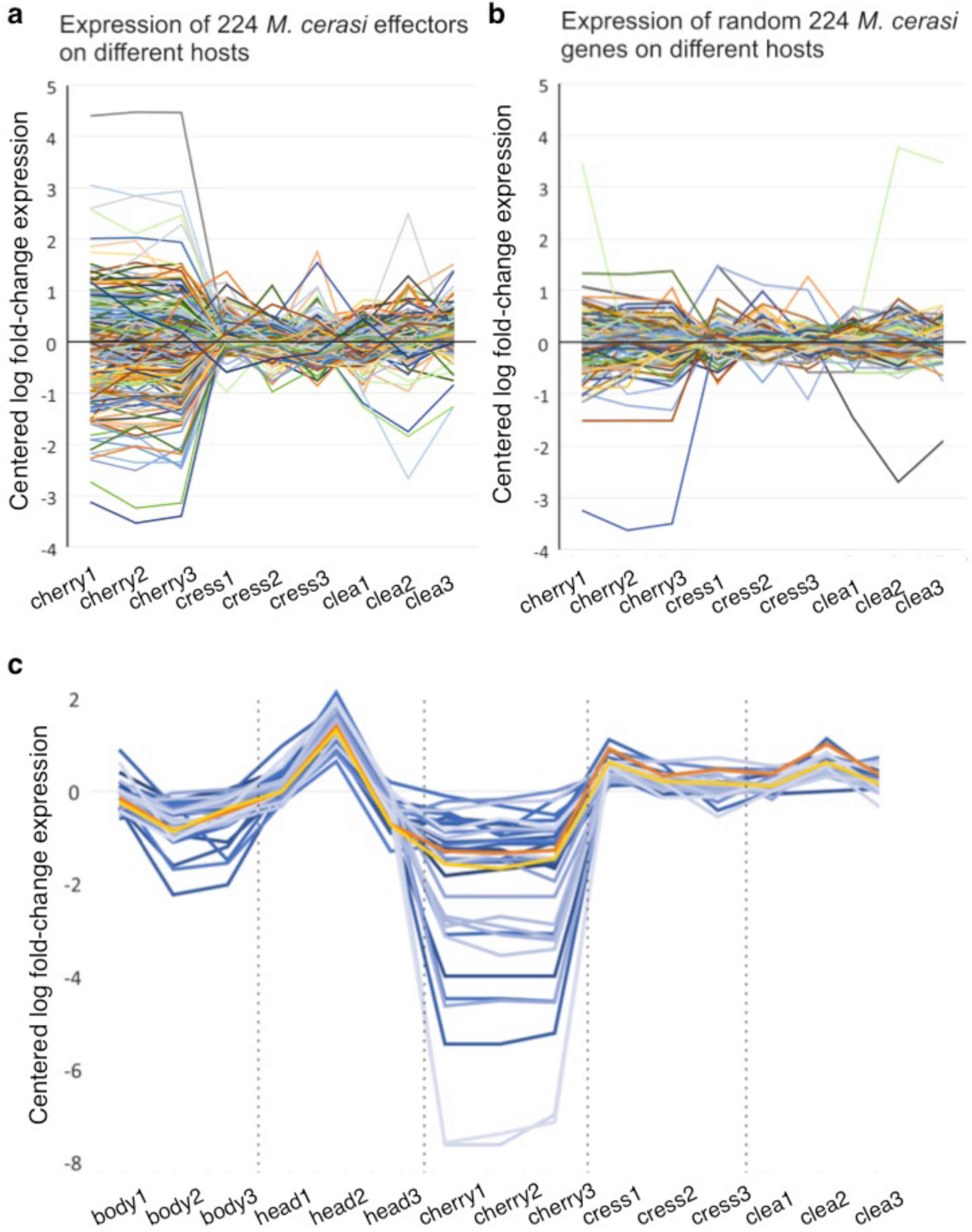
*Myzus cerasi* effector gene expression profiles across populations from different host environments. (A) Mean centered log fold-change expression of 224 *M. cerasi* putative effectors across aphid populations from different host environments, including primary host cherry, under field conditions, (cherry1-3), as well as cress (cress1-3) and cleavers (clea1-3), both in a controlled environment. (B) Mean centered log fold-change expression of 224 *M. cerasi* randomly selected genes across aphid populations from different host environments, including cherry (cherry1-3), cress (cress1-3) and cleavers (clea1-3). (C) Identification of all other genes in the *M. cerasi* genome that are co-regulated with the Mp1:Me10-like pair based on a >90% Pearson’s correlation across different populations (blue, n = 35). Mp1-like is indicated in orange and Me10-like in yellow.

### Break-down of aphid effector co-regulation in the aphid population collected from primary hosts under field conditions

We previously showed that expression of aphid effector genes, required for parasitism, was tightly co-regulated pointing to a mechanism of shared transcriptional control (Thorpe et al. 2018). To assess co-regulation in the different *M. cerasi* populations, we analyzed the co-expression patterns of Mc1 and Me10-like, an effector pair that is physically linked across aphid genomes and tightly co-regulated, together with all other genes. A total of 35 genes showed a high level of co-expression in the three different *M. cerasi* populations and across different aphid tissues (Fig. 4C). This number is much smaller compared to the set of co-regulated genes in *R. padi* (213) and *M. persicae* (114), which could be due to differences the quantity and quality of the RNAseq datasets we used for these analyses (Thorpe et al. 2018). However, the pattern of co-regulation observed in aphids collected from secondary host plants (controlled environment) is not apparent in aphids collected from the primary host cherry (field) (Fig. 4C), which affects the overall accuracy and ability to predict co-regulated genes. Possibly, the diversity of the *M. cerasi* population collected from primary hosts in the field underlies the observed break-down in co-regulation.

### Differential exon usage in *M. cerasi* populations

We also found evidence for differential exon usage when comparing the different aphid transcript datasets. Overall, 263 genes show significant differential exon usage when comparing aphid datasets associated with the different primary (field) and secondary hosts (controlled environment) (Table S7). These 263 genes contain 2551 exons, of which 443 show differential expression between aphid populations from primary hosts in the field versus populations from secondary hosts in a controlled environment. No significant GO annotation is associated with these 263 genes. One example of differential exon usage in *M. cerasi* is peroxidase gene Mca06436, which contains 5 exons, 2 of which are significantly more highly expressed in aphids collected from primary hosts in the field (Fig. 5). This suggests that alternative splicing may be associated with the adaptation of the field population to the secondary hosts in a controlled environment.

**Fig. 5.**
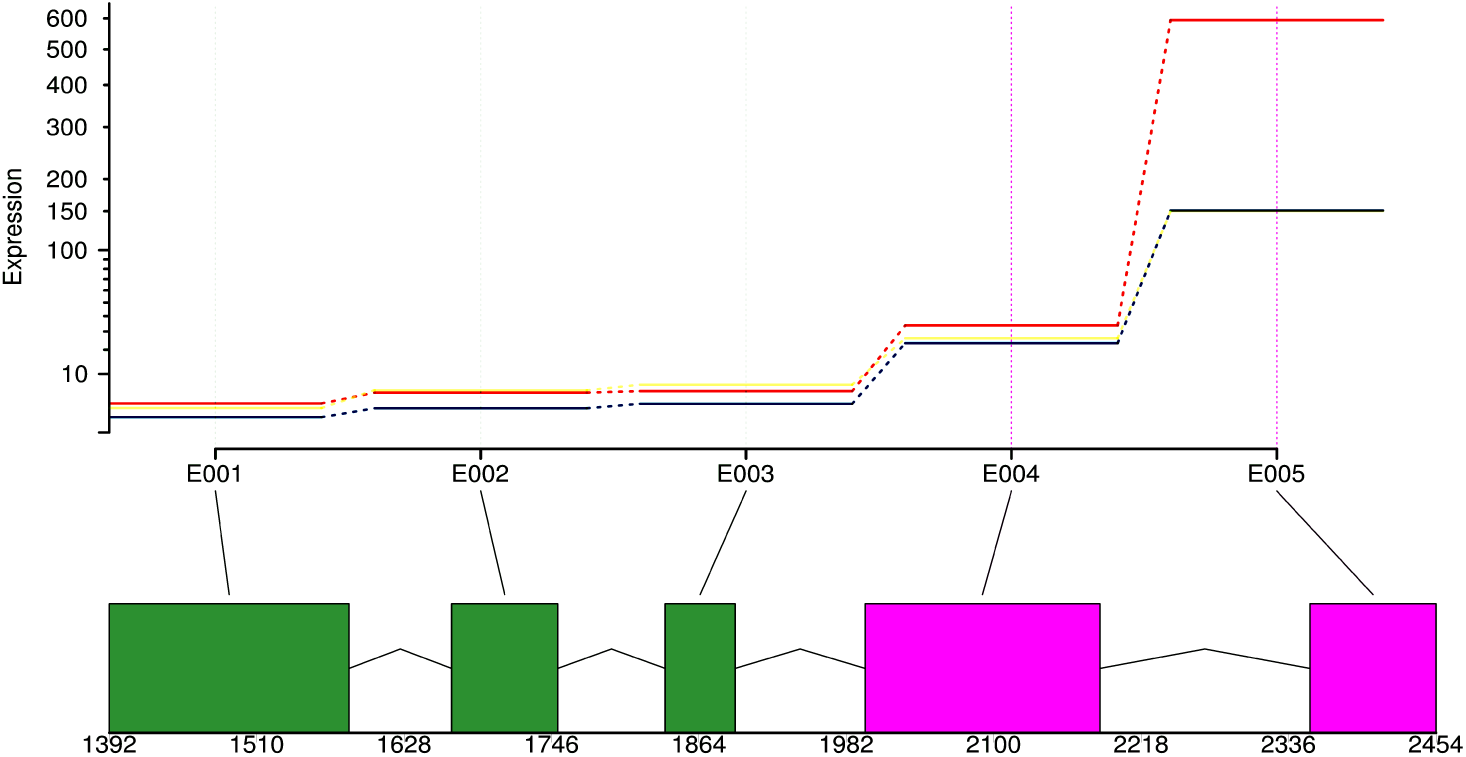
Graphical representation of differential exon usage observed in gene Mca06436|peroxidase-like in the transcriptome of *Myzus cerasi* populations from different host environments, including primary host cherry, in the field, (red line), as well as cress (yellow line) and cleavers (blue line), both in a controlled environment. The five different exons are indicated by E001-E005 and exons displaying significant differential expression are coloured pink. Numbers indicate nucleotide start and end positions of the different exons. The last exon shows 4 times greater expression in aphids collected from cherry in the field compared to those from cleavers or cress, both in a controlled environment.

### The majority of *M. cerasi* genes differentially expressed across different host environments are duplicated

Interestingly, the majority of genes differently expression across the three *M. cerasi* populations are duplicated (not single copy). For the genes upregulated in *M. cerasi* from cress (controlled environment) versus cherry (field) only 14% are single copy, which is significantly lower than the percentage of single copy genes in a randomly selected set of genes (p<0.001, Mann-Whitney U test). Moreover, for all sets of differentially expressed genes, the differentially expressed genes were more likely to be duplicated when compared to a background random gene set (p<0.001) (Table S8). To assess the categories of gene duplication within the differential expressed gene sets, 100 iterations of randomly selecting 100 genes were conducted to obtain a background population. This yielded a mean and standard deviation for each duplication type from the parent gene population (normally distributed). A probability calculator (Genstat) was used to determine how likely the observed counts were to occur at random. This showed that most of the duplicated differentially expressed genes were within the “dispersed duplication” category (p<0.001) and that there was no significant difference in the occurrence of tandem or proximal gene duplications (p>0.05) (Table S8). In contrast to predicted *M. cerasi* effectors, the differentially expressed genes identified in this study were not significantly further away from their neighbor in the 3’ direction (p=0.163, Mann-Whitney U Wilcoxon rank-sum test), or their 5’ neighbor gene (p=0.140, Mann-Whitney U Wilcoxon rank-sum test) when compared to an equal sized random population (Figure S3). Altogether our data suggest that *M. cerasi* multi-gene families may play an important role adaptation to host environments. This is in line with Mathers et al, (Mathers et al. 2017) who showed that duplicated genes play a role in adaptation of *M. persi*cae to different secondary host species.

## Conclusion

Aphids feature complex life cycles, which in some cases involve alternation between summer and winter host plant species. Here we show that, under controlled conditions, *M. cerasi* adaptation from primary to secondary host species does not readily occur, with only 10-20% aphid survival. Comprehensive gene expression analyses of *M. cerasi* populations collected from primary hosts in the field and adapted to secondary hosts under controlled conditions revealed sets of detoxification genes that are differentially regulated and differential exon usage associated with a change in host environment. Many of the differentially expressed genes are members of multi-gene families. In contrast, we find only limited transcriptional plasticity to secondary host switching under controlled conditions.

## Methods

### Aphid collection and adaptation

*M. cerasi* was collected in July 2013 from two separate locations in Dundee, United Kingdom. Mixed age aphids from branches of an infested cherry tree were flash frozen in liquid nitrogen upon collection (3 replicates of 50 aphids per location). For adaptation to secondary host plants, 50 aphids of mixed age, were transferred to *Galium aparine* (cleavers) or *Barbarae verna* (Land cress) detached branches placed in 3 replicate cup cultures per location. Aphid survival was assessed after 1 week. Then, 5 aphids of the surviving population on cleavers, were moved to a fresh cup culture containing detached cleavers branches. Fresh plant material was added to the cups after 2 weeks. One week later 50 mixed-age aphids per cup were flash frozen (aphids adapted to cleavers for RNAseq) and fresh cleavers branches together with Land cress branches were added to the cups. One week after adding the Land cress plant material, all cleavers material was removed and fresh cress branches were added and fresh plant material was regularly provided. Three weeks later 50 mixed-age aphids were collected per cup culture and flash frozen (aphids adapted to cress for RNAseq). Aphids were maintained in cup cultures in controlled environment cabinets at 18°C with a 16 hour light and 8 hour dark period.

### RNA sample preparation and sequencing

Aphid samples were ground to a fine powder and total RNA was extracted using a plant RNA extraction kit (Sigma-Aldrich), following the manufacturer’s instructions. We prepared three biological replicates for *M. cerasi* collected from each host. RNA quality was assessed using a Bioanalyzer (Agilent Technologies) and a Nanodrop (Thermo Scientific). RNA sequencing libraries were constructed with an insert size of 250bp according to the TruSeq RNA protocol (Illumina), and sequenced at the previous Genome Sequencing Unit at the University of Dundee using Illumina-HiSeq 100bp paired end sequencing. All raw data are available under accession number PRJEB24338.

### Quality control, RNAseq assembly and differential expression

The raw reads were assessed for quality before and after trimming using FastQC (Andrews 2010). Raw reads were quality trimmed using Trimmomatic (Q22) (Bolger and Giorgi), then assembled using genome-guided Trinity (version r20140717) (Grabherr et al. 2011). Transrate was run twice to filter out low supported transcripts (Smith-Unna et al. 2015).

RNAseq assembly and annotation is available at DOI: 10.5281/zenodo.1254453. For differential gene expression, reads were mapped to the *Myzus cerasi* genome (Thorpe et al. 2018), per condition using STAR (Dobin et al. 2013). Gene counts were generated using Bedtools (Quinlan and Hall 2010). Differential gene expression analysis was performed using EdgeR (Robinson et al. 2010), using LOG fold change >2, FDR p<0.001 threshold. GO enrichment analysis was performed using BLAST2GO (version 2.8, database September 2015) (Conesa et al. 2005) using FDR 0.05. The genome annotations were formatted using GenomeTools (Gremme et al. 2013) and subsequently HTseq (Anders et al. 2015) was used to quantify exon usage. Differential exon expression was performed using DEXSEQ FDR p<0.001 (Anders et al. 2012). Heatmaps were drawn as described in Thorpe et al. (Thorpe et al. 2018).

Gene duplication categories were used from Thorpe et al. (Thorpe et al. 2018). From these data, a random population was generated by running one hundred iterations on a set of 100 randomly selected genes and their duplication types for subsequence statistical analyses. The script to generate random mean and standard deviation counts is available on Github (https://github.com/peterthorpe5/Myzus.cerasi_hosts.methods). Statistical analysis was performed using Probability Calculator in Genstat (17^th^ edition). The obtained value from the gene set of interest (differentially express genes across aphid populations) was compared to the distribution of the random test set. Datasets identified as being significantly different from the random population did not significantly deviate from a normal distribution, thus the data was normally distributed. To assess the distances from one gene to the next, an equal sized population (1020) of random genes and their values for distance to their neighboring gene in a 3’ and 5’ direction was generated. The real value and random values were not normally distributed and were analyzed in Genstat (17^th^ edition) using a non-parametric Mann-Whitney U Wilcoxon rank-sum test.

For SNP identification, RNAseq data was mapped back to the reference genome using STAR (2.5.1b) with --outSAMmapqUnique 255, to allow only unique mapping (Dobin et al. 2013). SNPs were identified using Freebayes (Garrison and Marth 2012). VCFtools (0.1.15) (Danecek et al. 2011) was used on the resulting vcf files to identify SNPs per 10Kb.

### Validation of expression profiles by qRT-PCR

Validation of the RNA-Seq experiment was completed with the Universal Probe Library (UPL) RT-qPCR system (Roche Diagnostics ©). RNA samples analyzed were those used for RNAseq analyses (aphid collections from location 1) as well as samples from aphids collected at a separate location (location 2) and adapted to secondary hosts (3 biological replicates). For all experiments, aphid RNA was extracted using RNeasy Plant Mini Kit (Qiagen). RNA samples were DNAse treated with Ambion® TURBO DNA-free™. SuperScript® III Reverse Transcriptase (Invitrogen) and random primers were used to prepare cDNA. Primers and probes were designed using the predicted genes sequences generated in the RNA-seq data analysis and the Assay Design Center from Roche, selecting “Other organism” (https://lifescience.roche.com/en_gb/brands/universal-probe-library.html). Primers were computationally checked to assess if they would amplify one single product using Emboss PrimerSearch. Primers and probes were validated for efficiency (86-108 %) before gene expression quantification; five dilutions of threefolds for each primer pair-probe were used for generating the standard curve. The 1:10 dilution of cDNA was selected as optimal for RT-qPCR using the UPL system. Reactions were prepared using 25µl of total volume, 12.5µl of FastStart TaqMan Probe Master Mix (containing ROX reference dye), 0.25µl of gene-specific primers (0.2mM) and probes (0.1mM). Step-One thermocycler (Applied Biosystems by Life Technology©) was set up as follows: 10 min of denaturation at 95°C, followed by 40 cycles of 15 s at 94°C and 60 s at 60°C. Relative expression was calculated with the method ΔCt (Delta Cycle threshold) with primer efficiency consideration. Three technical replicates were run per sample. Reference genes for normalization of the cycle threshold values were selected base on constant expression across different conditions in the RNA-Seq experiment. The reference genes were CDC42-Kinase (Mca01274), actin (Mca10020) and tubulin (Mca04511). The fold change calculations were done by ΔΔ Ct method (Delta Cycle threshold) and primer efficiency was taken into consideration.

## Acknowledgement

We thank Brian Fenton for help with adapting *Myzus cerasi* collected from cherry trees to secondary hosts, and Melanie Febrer for advice on the RNAseq experiment and sample processing.

## Availability of data and materials

All data are available under accession numbers PRJEB24338. *Myzus cerasi* genome and annotation was downloaded from http://bipaa.genouest.org/is/aphidbase/ and DOI:10.5281/zenodo.1252934. All custom python scripts used to analyse the data use Biopython (Cock et al. 2009), as well as details on how they where applied for data analyses are available on https://github.com/peterthorpe5/Myzus.cerasi_hosts.methods, and DOI: 10.5281/zenodo.1254453.

## Authors’ contributions

PT and JIBB conceived the experiments, PT and CEM performed sample preparation and qRT-PCR analyses, PT, CEM, SEvdA and JIBB analyzed data, PT and JB wrote the manuscript, with input from all authors. All authors read and approved the final manuscript.

## Supplementary data

**Fig. S1.**
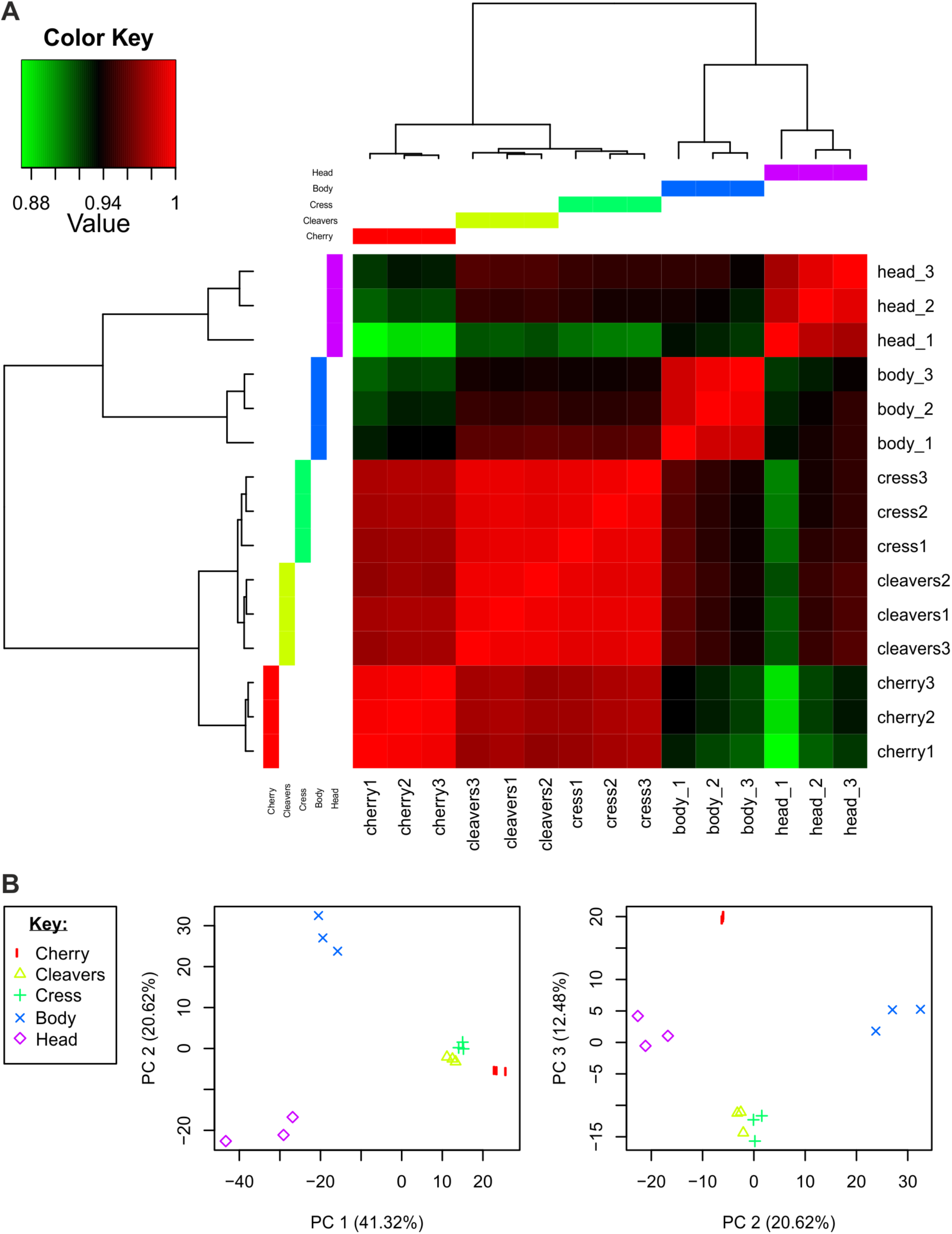
Transcriptome differences between *Myzus cerasi* populations from different host environments Genome-wide analysis of *M. cerasi* transcriptional responses to interaction primary host cherry (field) or secondary hosts cress and cleavers (both in controlled environment), and comparison to previously published tissue-specific transcriptome of dissected heads and bodies (Thorpe et al. 2016). (A) Clustering of transcriptional responses reveals that *M. cerasi* gene expression is different in populations from the different host environments and also that expression in head and body tissues can be separated based on these analyses (B). Principle component analysis. The top 3 most informative principle components describe approximately 75% of the variation, and separate the both the host species interaction data as well as tissue-specific data well.

**Table S1.** List of 934 differentially *Myzus cerasi* genes across different host environments.

**Table S2.** List of significant GO-terms associated with genes differentially expressed in cluster A and E (Fig. 2).

**Table S3.** List of significant GO-terms associated with genes differentially expressed across different *Myzus cerasi* host environments, corresponding to Fig. 3A.

**Table S4.** List of significant GO-terms associated with genes differentially expressed across different *Myzus cerasi* host environments, corresponding to Fig. 3B.

**Table S5.** List of significant GO-terms associated with genes differentially expressed across different *Myzus cerasi* host environments, corresponding to Fig. 3C.

**Fig. S2.**
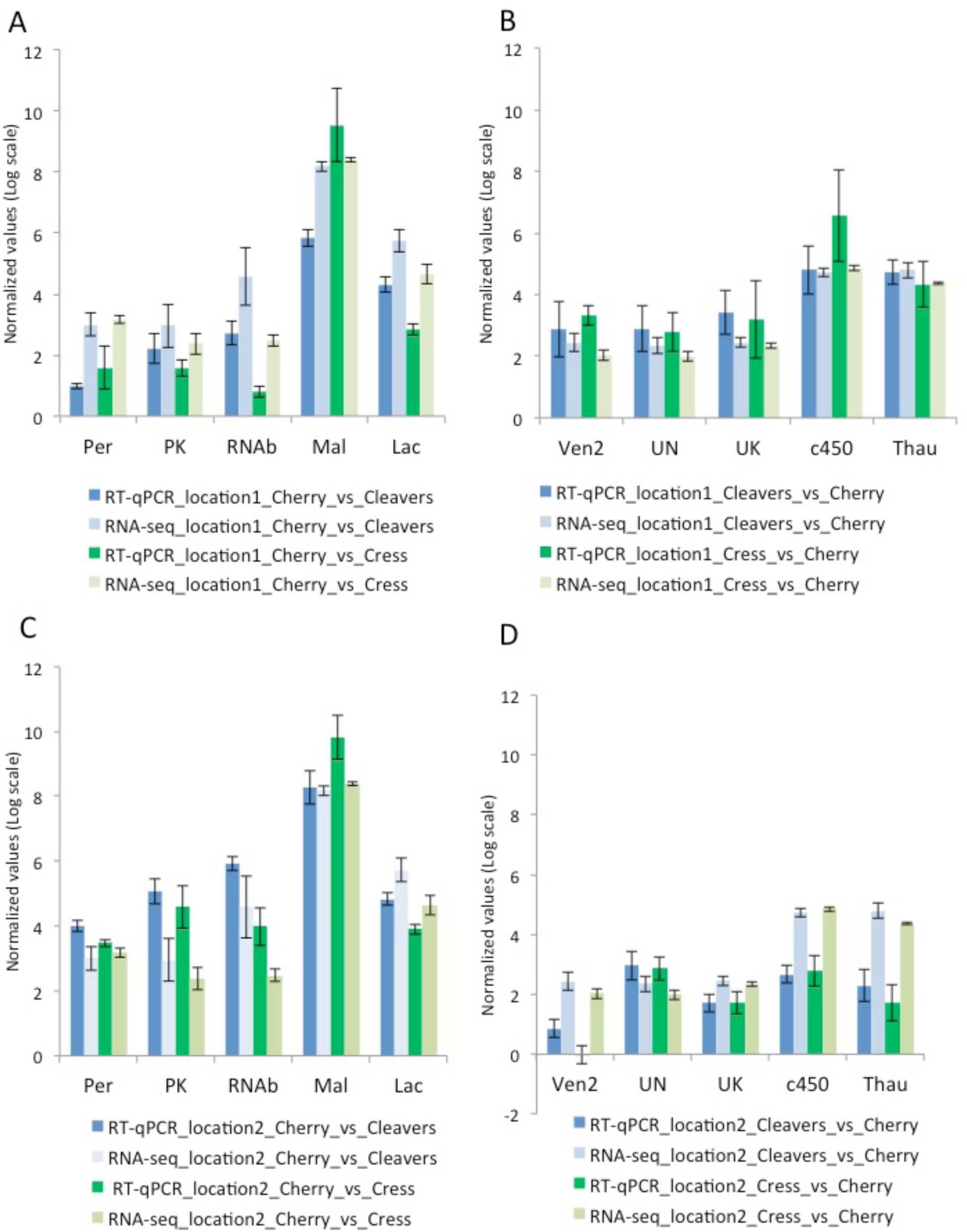
Validation of differential gene expression by qRT-PCR (A) Genes up-regulated during cherry (field) versus cleavers/cress (controlled environment) interactions in the *Myzus cerasi* population collected from location 1. (B) Genes up-regulated during the cleavers/cress (controlled environment) versus the cherry (field) interactions in the *M. cerasi* population collected from location 1. (C) Genes up-regulated during cherry (field) versus cleavers/cress (controlled environment) interactions in the *M. cerasi* population collected from location 2. (D) Genes up-regulated during the cleavers/cress (controlled environment) versus the cherry (field) interactions in the *M. cerasi* population collected from location 2. The validated genes up-regulated during the cherry interactions were peroxidase (Mca14094-Per), protein kinase (Mca07516-PK), RNA binding (Mca07514-RNAb), maltase (Mca25862-Mal) and lactase (Mca19306-Lac). Validated genes up-regulated during the cress/cleavers interactions were venom protein (Mca05785-Ven), uncharacterized protein (Mca06816-UN), unknown protein (Mca06864-UK), cytochrome 450 (Mca22662-c450) and thaumatin (Mca12232-Thau). Blue and green series represent RT-qPCR validation results and pale blue and pale green represent RNA-seq results. Error bars indicate standard error.

**Table S6.** List of 224 *Myzus cerasi* putative effectors and their expression levels across different host environments.

**Table S7.** List of *Myzus cerasi* genes showing differential exon usage across different host environments.

**Table S8.** Gene duplication types in *Myzus cerasi* genes differentially expressed across different host environments.

**Figure S3.**
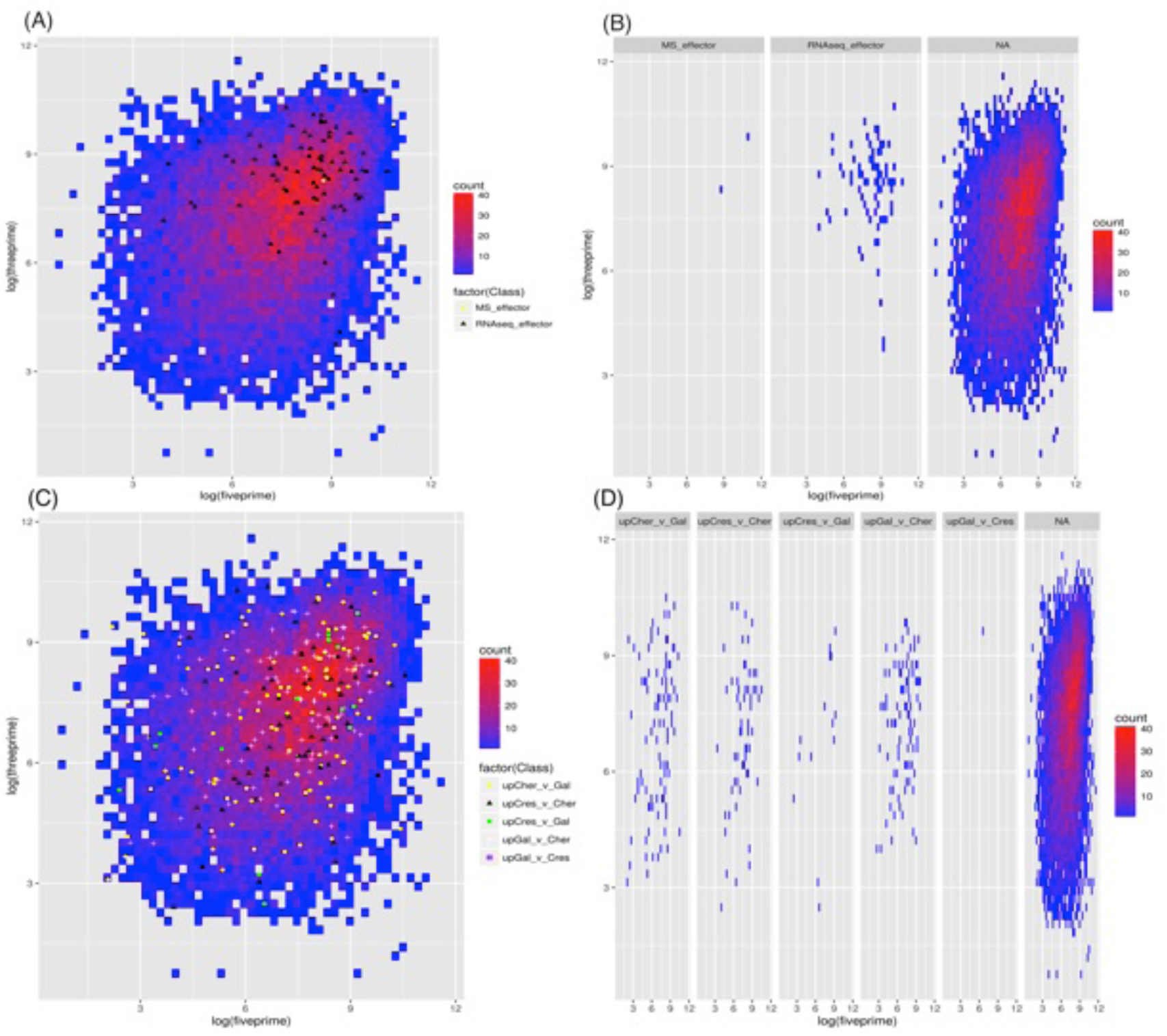
Heat maps graphically representing the LOG nucleotide distance from one gene to its neighboring genes in a 3’- and 5’ -direction. Various gene categories are colored and coded in the relevant keys. (A) and (B) Geneic distance heat map for predicted effectors, which were significantly further away from their neighboring genes and thus in gene sparse regions (Thorpe et al. 2018). (C) and (D) Genic distances for differentially expressed genes identified in this study. These are not significantly further away from their neighboring genes in either 3’- or 5’-direction.

