## Supplementary figures and images for "Extensive transcriptional changes in the aphid species *Myzus cerasi* under different host environments associated with detoxification genes"

### Figure S1

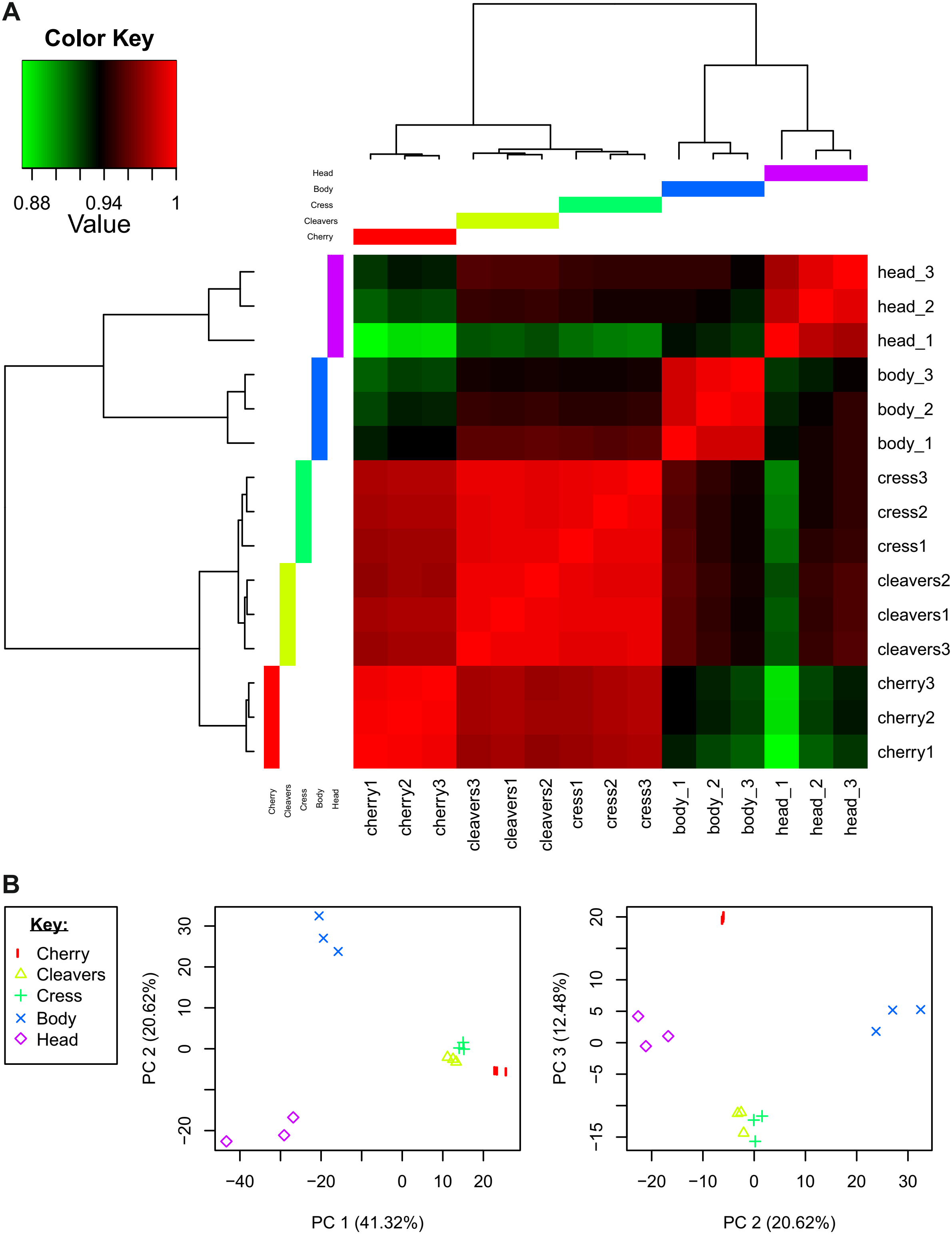

### Figure S2

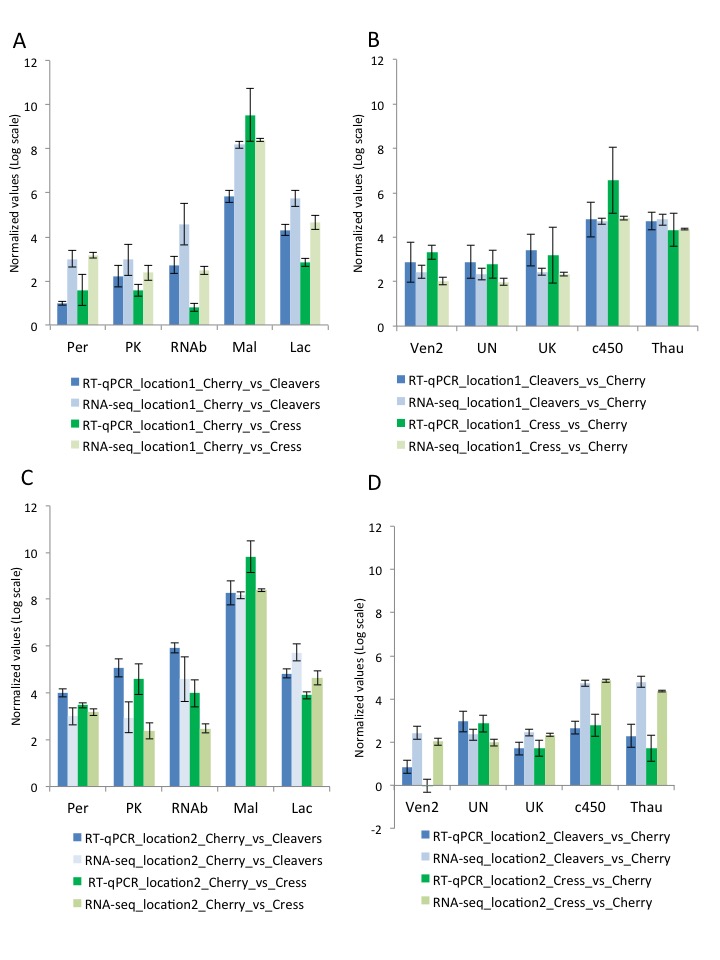

### Figure S3

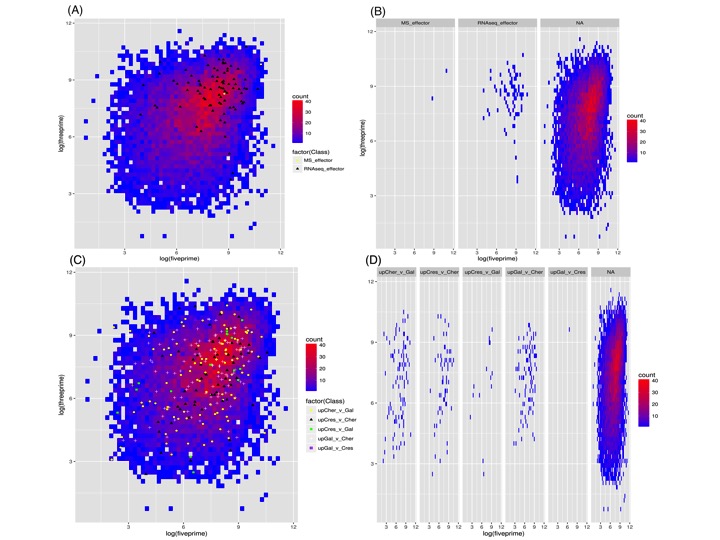
